# Mining novel cis-regulatory elements from the emergent host *Rhodosporidium toruloides* using transcriptomic data

**DOI:** 10.1101/2022.10.03.510576

**Authors:** Luísa Czamanski Nora, Murilo Henrique Anzolini Cassiano, Ítalo Paulino Santana, María-Eugenia Guazzaroni, Rafael Silva-Rocha, Ricardo Roberto da Silva

## Abstract

The demand for robust microbial cell factories that can produce valuable biomaterials while being resistant to stresses imposed by current bioprocesses is rapidly growing. *R. toruloides* is an emerging host that presents desirable features for bioproduction, since it can grow in a wide range of substrates and tolerate a variety of toxic compounds. In order to explore *R. toruloides* suitability for application as a cell factory in biorefineries, we sought to understand the transcriptional responses of this yeast when growing under experimental settings that simulated those used in biofuels-related industries. Thus, we performed RNA sequencing of the oleaginous, carotenogenic yeast in different contexts. The first ones were stress-related: two conditions of high temperature (37 °C and 42 °C) and two ethanol concentrations (2% and 4%), while the other was using the inexpensive and abundant sugarcane juice as substrate. Using transcriptomic data, differential expression and functional analysis were implemented to select differentially expressed genes and enriched pathways from each set-up. A reproducible bioinformatics workflow was developed for mining new regulatory elements. We then predicted, for the first time in this yeast, binding motifs for several transcription factors, including HAC1, ARG80, RPN4, ADR1, and DAL81. Most of the putative transcription factors uncovered here were involved in stress responses and found in the yeast genome. Our method for motif discovery provides a new realm of possibilities in the study of gene regulatory networks, not only for the emerging host *R. toruloides*, but for other organisms of biotechnological importance.

## 1. Introduction

Microbial cell factories are in ever-growing demand due to the pursuit of more sustainable products that support the concepts of green chemistry and biorefineries (Cherubini, 2010; Martins *et al*., 2021). The benefits go beyond obtaining valuable products from low-cost raw materials, but include the independence from oil, a finite and extremely polluting material (Borodina and Nielsen, 2014). Innovations on obtaining products from renewable substrates are based on the genetic improvement and metabolic engineering of microorganisms that are able to degrade and ferment different biomasses (Martins-Santana *et al*., 2018). The main challenges the currently available cell factories face is not only the capability to break complex compounds into simpler chains, but also to be able to tolerate adverse conditions resulting from bioprocesses (Fletcher *et al*., 2017).

*Rhodosporidium toruloides* is emerging as a potential, robust host for the completion of both tasks. This oleaginous, carotenogenic yeast can flourish on a wide range of substrates, such as sugarcane juice and bagasse, crude glycerol, wheat straw, cassava starch, among others (Bonturi *et al*., 2017; Park, Nicaud and Ledesma-Amaro, 2018; Lopes, 2020). It can also store high amounts of lipids – up to 60% of its total cell weight – and tolerate compounds commonly toxic for other yeast species (Saini *et al*., 2020). It performs better than the widely used baker’s yeast, *Saccharomyces cerevisiae*, for example, as it can degrade C5 sugars and lignin-derived compounds, which the latter cannot (Nora, Westmann, *et al*., 2019).The advantages go over other lipid-producing strains as well, such as *Yarrowia lipolytica*, since it can grow in more complex carbon sources and presents higher tolerance to several inhibitors like 5-hydroxy methyl ester, furfural, acetic acid and vanillin (Saini *et al*., 2020). *R. toruloides* has already been shown to produce several different bioproducts in a variety of conditions. Examples of these bioproducts include enzymes, carotenoids and lipid-based compounds: linoleic acid, oleic acid and cocoa butter substitute (Park, Nicaud and Ledesma-Amaro, 2018). Recently, terpenes that can be used as precursors of biodiesel and jet fuel alternatives were produced in pilot scale, derived from corn stover hydrolysates (Yaegashi *et al*., 2017; Kirby *et al*., 2021). Hence, all these distinctive characteristics point to *R. toruloides* as a great candidate to improve the productivity of biorefineries.

There is great interest in using unconventional, more robust yeasts in detriment to the use of *S. cerevisiae* for ethanol production in bioreactors (Wehrs *et al*., 2019; Wu *et al*., 2020). Ethanol tolerance and thermotolerance, for instance, are desirable features. First, ethanol accumulation during fermentation can become a significant stress factor during the process (Stanley *et al*., 2010). Secondly, higher fermentation temperatures can lower the cost of cooling bioreactors during the process and reduce bacterial contamination, decreasing the need for antibiotics (Abdel-Banat *et al*., 2010; Huang *et al*., 2018). Thus, in order to explore if *R. toruloides* could tolerate and thrive in these stressful environments, we decided to investigate the transcriptional response of this yeast growing in high temperatures and in the presence of ethanol. Furthermore, Soccol *et al*. (2017), developed a process of producing biofuels using *Rhodosporidium* grown in a simple media containing only sugarcane juice and urea. Sugarcane juice is an abundant and low-cost substrate, containing up to 15% of fermentable sugars. Based on their work, we additionally sought to understand the changes in transcription of *R. toruloides* grown in media containing sugarcane juice and urea (Soccol *et al*., 2017).

Transcription factors (TF) are proteins that bind to specific DNA sequences modulating the rate of transcription of DNA to mRNA. These specific DNA sequences are collectively called transcription factor binding sites (TFBS), or binding motifs, and are present in the *cis*-regulatory region of DNA surrounding the transcription starting sites. The activity of TFs can act directly or indirectly to transform physiological and environmental signals into patterns of gene expression. Thus, identifying both the proteins and the position of their binding motifs is essential for understanding gene regulation in the cell and allowing better transcription control, which can serve as a basis for new bioengineering applications (Martins-Santana et al., 2018). However, finding new regulatory sequence motifs is not an easy task, especially in eukaryotic organisms. One of the reasons is that primary nucleotide sequence is not the only characteristic that specifies a target. Additional factors, such as genomic context, DNA binding, DNA modification and DNA shape can change nucleotide preferences (Inukai, Kock and Bulyk, 2017). Besides, it is impossible to test all the environmental conditions that a natural regulatory network is able to respond to. Nevertheless, some tools were developed aiming to predict regulatory elements in specific datasets, helping to overcome limitations in motif discovery.

Here, we developed a new pipeline for discovery of *cis*-regulatory elements integrating tools for motif prediction and applied it to the emergent microbial cell factory *R. toruloides*, which can later be expanded to other biotechnologically relevant organisms. In order to do that, we cultivated *R. toruloides* in the settings described above and extracted RNA samples from early time points of growth. Then, we performed RNA sequencing, functional analysis of the transcriptional responses and discovery of new motifs from genes of the main pathways found to be enriched in each specific condition. In this sense, we are providing new insights regarding the regulation of *R. toruloides* genes in a transcriptomic approach and describing, for the first time, putative TFBS that can be used for transcriptional control in this yeast, offering the basis for building new genetic tools for this promising host.

## 2. Material and Methods

### 2.1. Strains, media and growth conditions

For RNA sequencing experiments using *R. toruloides* grown in a medium with sugarcane juice (SCJ), cultures were first grown in LB medium (1% yeast extract, 1% tryptone and 0.5% NaCl) for 24 hours. LB medium was used in the inoculum so that there was no other sugar influencing the yeast metabolism. The inoculum was centrifuged and transferred to media containing sugarcane juice. Considering the concentration of 120g of sucrose per liter of juice, we standardized the final concentration of the media as 4% sucrose. In addition, 1% urea was added to be used as a nitrogen source. The filtered sugarcane juice was pasteurized at 65°C for 30 minutes before being added to the autoclaved medium. SCJ cultures were grown for 8 hours and then collected for RNA extraction. Aliquots of the inoculum in LB grown for 24 hours were also collected to serve as internal control and henceforth will be called time 0 h. The 8-hour growth time was selected for this condition because the purpose was to analyze the genes as soon as there was expression of the genes for adaptation to the new medium.

The experiments for the stress conditions were carried out as follows: the inoculum was grown in YPD medium (1% yeast extract, 2% peptone and 2% glucose) at 200 rpm at 30 °C for 24 hours. These inoculants were centrifuged and transferred to the respective media, containing YPD and one of the respective stress conditions: 2% ethanol and 4% ethanol, which were grown in a shaker at 30 °C, and the high temperature conditions that were grown at 37 °C and 42 °C. The control condition was determined as the inoculum grown in YPD for 24 hours. After being transferred to the respective media, the cultures were grown for another 16 hours and collected for RNA extraction and sequencing. All inoculations were performed in a 1:10 ratio, and all experiments were performed in triplicates. The strain *R. toruloides* IFO0880 was used in all experiments (Lin *et al*., 2014; Coradetti *et al*., 2018).

### 2.2. RNA extraction and sequencing

Cell lysis was performed by pelleting yeast cells from the culture by centrifugation and resuspending it with Trizol^®^ (Thermo Fisher Scientific). Samples were transferred to tubes containing zirconium beads and lysed using a cell homogenizer. RNA extraction using Trizol^®^ (Thermo Fisher Scientific) was performed following the manufacturer’s instructions. The resulting RNA samples were immediately purified using the Qiagen^®^ Rneasy mini kit. Samples were submitted to analysis by Agilent Bioanalyzer RNA 6000 Nano kit for RNA integrity check, aiming for a RIN number higher than 8. After being extracted and purified, all RNA samples were kept in a −80 °C freezer. The samples were sent for sequencing at the Genomics Center of the University of São Paulo in Piracicaba, São Paulo, Brazil. The sequencing library was prepared by the facility using Illumina TruSeq Stranded mRNA Sample Prep LT Protocol. RNA sequencing was performed using the HiSeq SBS v4 kit in Illumina HiSeq 2500, with paired reads of 100 bp (2×101). Raw data is available as Sequencing Read Archives (SRA) on the NCBI website under accession number PRJNA883675.

### 2.3. RNA-seq differential gene expression analysis

Read quality check was performed using FastQC (Andrews, 2018) and the trimming was made using the Trimmomatic software (Bolger, Lohse and Usadel, 2014). Henceforth, we used only the reads assigned by Trimmomatic as paired. Gene expression was quantified with kallisto (Bray *et al*., 2016) using for counting the *R. toruloides* IFO0880 JGI reference transcriptome (Coradetti *et al*., 2018) available at https://mycocosm.jgi.doe.gov/Rhoto_IFO0880_4/Rhoto_IFO0880_4.home.html. The differential expression analysis was done using DESeq2 (Love, Huber and Anders, 2014) in the R platform (R Core Team, 2019). Genes whose adjusted *p-*valuesn (*adjpval*) were lower than 0.05 and whose log2-fold change values were lower than −1 or higher than 1 were selected as differentially expressed genes (DEGs).

### 2.4. Functional and pathway analysis

In order to analyze the DEGs function, we used the reference gene function annotation file created by KOG (also provided by JGI, as above). For pathway enrichment analysis we used the GAGE R package (Luo *et al*., 2009). As input in this step, we used the DEGs from each condition with their function annotated by the KEGG Automatic Annotation Server (https://www.genome.jp/tools/kaas/), as this tool provides a compatible file for GAGE analysis. The enriched pathways were filtered by a p-value lower than 0.05. Bubble maps were plotted using R package ggplot2 (Wickham, 2016). Pathview R package (Luo and Brouwer, 2013) was used to visualize enriched pathways whenever needed.

### 2.5. Putative *cis*-regulatory elements discovery

For each condition, we selected the DEGs that belong to the KOG classes. We extracted the promoter sequence of these genes on the reference database and applied the HOMER software (Heinz *et al*., 2010) in each promoter sequence sets, with the following parameters: -len 10, -stat hypergeo, -olen 3, -strand both, - maxBack 0.3. The background file was the *R. toruloides* IFO0880 JGI reference promoter sequences. We performed a pairwise comparison between all motifs, by applying the universalmotif (Tremblay, 2020) function ‘compare_motifs’ using Pearson Correlation Coefficient (PCC) as method and average mean as scoring strategy. We also performed pairwise alignment using TOMTOM, from MEME suite (Gupta *et al*., 2007), providing as input our motif file as both target and source, with the following arguments: -min-overlap 0, -evalue, -thresh 10.0 and -dist pearson. Finally, we filtered the results by a p-value lower than 0.05, an e-value lower than 1 and a PCC higher than 0.8. All motifs from our experimental settings were found using HOMER and were aligned with the ones from JASPAR2018_CORE_fungi_non-redundant database using TOMTOM. Subsequently, they were plotted and analyzed as a graph, using Gephi (Bastian, Heymann and Jacomy, 2009), applying Circle Pack Layout for representation with the variables ‘predicted or annotated’, experimental condition and node degree as hierarchy.

### 2.6. Finding TF candidates

For the overrepresented motifs found on JASPAR alignment, we got the amino acid sequence of the corresponding transcription factor protein and then performed a blast on JGI platform, to search for similar proteins on the *R. toruloides* reference assembled genome (provided by JGI, as above). Amino acid sequences and GO functions of each predictive TF were extracted from UniProt (http://uniprot.org).

### 2.7. Bioinformatics Workflow

All analyses performed on the R environment were organized in an easy-to-use Jupyter Notebook, the customized code and instructions to install and run the workflow are publicly available at https://github.com/computational-chemical-biology/cis_reg.

## 3. Results

### 3.1. A new pipeline for discovery of regulatory elements from transcriptomic data

**Figure 1** summarizes the workflow developed in this work. The experimental design consisted of growing cells and extracting their RNA as described in the Methods section. Then, the RNA sequencing data was used as input for the motif discovery pipeline. First, differential expression analysis was performed in order to select the Differentially Expressed Genes (DEGs), and functional analysis was executed to find enriched pathways for each condition. We then extracted the promoter sequences from the genes of the specific enriched pathways and applied the HOMER software for motif prediction. **Table 1** contains the number of promoter sequences and motifs found by HOMER for each case. We performed an internal pairwise comparison between all motifs, by applying both the universalmotif package and TOMTOM tool. Subsequently, we performed an external comparison against the Jaspar database using TOMTOM. The entire workflow will be described in detail throughout the Results section.

**Figure 1.**
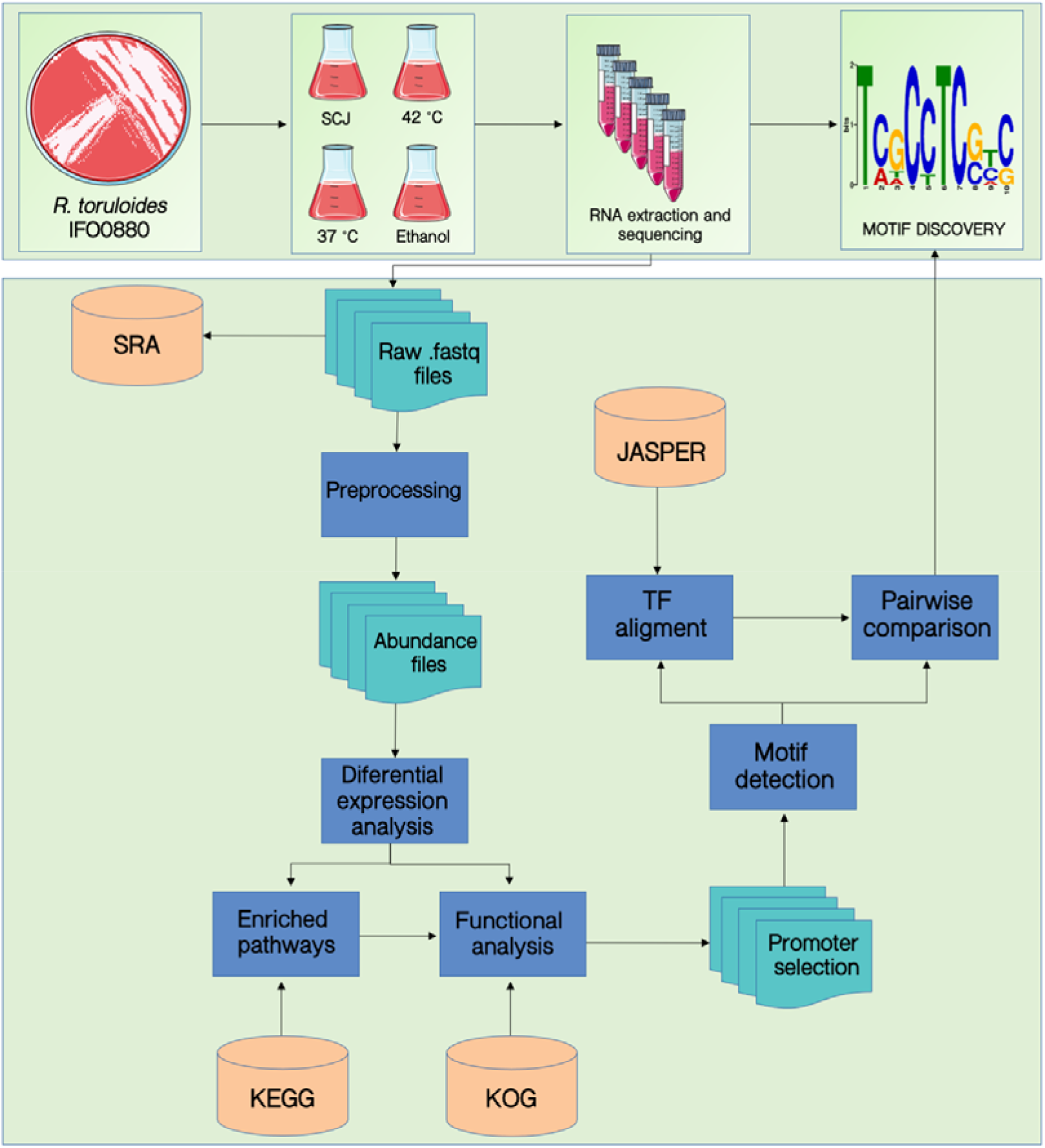
Workflow for the discovery of *cis*-regulatory elements using transcriptomic data.

**Table 1.** Mining of promoter and TFBS sequences. Number of promoter sequences selected for each condition and each KOG class and the number of possible motifs found by HOMER.

| Condition | KOG Class | Promoters | Motifs |
| --- | --- | --- | --- |
| **SCJ** | Amino acid transport and metabolism | 111 | 5 |
|  | Carbohydrate transport and metabolism | 105 | 5 |
|  | Chromatin Structure and dynamics | 39 | 16 |
|  | Secondary metabolites biosynthesis, transport and catabolism | 59 | 4 |
|  | Signal Transduction Mechanisms | 134 | 4 |
|  | Transcription | 76 | 5 |
|  | Translation, ribosomal structure and biogenesis | 140 | 5 |
| **Temperature 37°C** | Amino acid transport and metabolism | 20 | 2 |
|  | Carbohydrate transport and metabolism | 23 | 3 |
|  | Chromatin Structure and dynamics | 8 | 5 |
|  | Secondary metabolites biosynthesis, transport and catabolism | 23 | 3 |
|  | Signal Transduction Mechanisms | 23 | 2 |
|  | Transcription | 6 | 5 |
|  | Translation, ribosomal structure and biogenesis | 8 | 5 |
| **Temperature 42°C** | Amino acid transport and metabolism | 97 | 4 |
|  | Carbohydrate transport and metabolism | 106 | 5 |
|  | Chromatin Structure and dynamics | 26 | 5 |
|  | Secondary metabolites biosynthesis, transport and catabolism | 76 | 5 |
|  | Signal Transduction Mechanisms | 96 | 3 |
|  | Transcription | 60 | 5 |
|  | Translation, ribosomal structure and biogenesis | 93 | 5 |
| **Ethanol 2%** | Amino acid transport and metabolism | 37 | 4 |
|  | Carbohydrate transport and metabolism | 32 | 0 |
|  | Chromatin Structure and dynamics | 18 | 5 |
|  | Secondary metabolites biosynthesis, transport and catabolism | 35 | 5 |
|  | Signal Transduction Mechanisms | 30 | 2 |
|  | Transcription | 9 | 0 |
|  | Translation, ribosomal structure and biogenesis | 15 | 0 |
| **Ethanol 4%** | Amino acid transport and metabolism | 39 | 5 |
|  | Carbohydrate transport and metabolism | 39 | 5 |
|  | Chromatin Structure and dynamics | 10 | 0 |
|  | Secondary metabolites biosynthesis, transport and catabolism | 34 | 4 |
|  | Signal Transduction Mechanisms | 39 | 4 |
|  | Transcription | 7 | 0 |
|  | Translation, ribosomal structure and biogenesis | 17 | 0 |

### 3.2. Differential expression analysis shows a significant transcriptional change in sugarcane and high temperature conditions

The RNA sequencing data were analyzed through the RNA-seq differential analysis using DESeq2. For all analyses, there were two different condition settings: the first one was the sugarcane juice condition (SCJ) and the second one was the stress conditions (comprising 37 and 42 °C and ethanol 2% and 4%). The differential expression of *R. toruloides* genes grown on the SCJ condition was measured relative to its own control, which was the inoculum grown in LB for 24 hours (also called time 0h). Yet, the differential expression of genes from *R. toruloides* grown in stress conditions were measured using the inoculum grown in YPD for 24 hours. Each experimental setting was always analyzed separately since they had different internal controls. The principal component analysis (PCA) was performed to check the quality of the sequencing samples, where we could observe the consistency of our sample replicates in a summarized representation. As shown in the PCA graphs in **Supplementary Figure S1A**, where the two first components explain 84% of the variance, the replicates of each condition are close to each other and distant from their respective controls, which indicates that the gene expression differences are consistent inside each condition, allowing the comparison between them. Nevertheless, in **Supplementary Figure S1B**, we can see that the replicates of the culture grown at 37 °C are very close to the control condition. While the 42 °C is separated by the first principal component, explaining alone 50.1% of the variance.

To further investigate the quality of the RNA-seq experiments, the distance between samples, based on the transcript expressions, was measured (**Supplementary Figure S2**). In the heatmap analysis, Euclidean distance was applied to the expression pattern of each sample, where darker blue means more proximity and lighter blue means greater distance between samples. We can see the same trend in which the replicates of each condition are grouped, which confirms the PCA analysis. Similar to PCA, in **Supplementary Figure S2A**, we can see that the ethanol 2% and ethanol 4% conditions were not distant from each other. This shows that the changes in transcriptional behavior in *R. toruloides* were not divergent between the two conditions, albeit they are both distant from the control. In **Supplementary Figure S2B** it is possible to see a similar trend to PCA, where the SCJ condition is distant from its internal control.

The Venn Diagram depicted in **Figure 2** summarizes the DEGs found for *R. toruloides* grown in each experimental condition. The conditions in which more genes presented transcriptional change were the SCJ and the temperature 42 °C. More than three thousand DEGs were found for those experimental settings, while there were around one thousand DEGs for each of the other conditions: 37 °C and ethanol 2% and 4%. These results show that the latter conditions, although causing a significant change in the transcriptional behavior of *R. toruloides*, did not cause as much impact as the change to a rich substrate caused by the addition of sugarcane in the media nor the increase in temperature by growing at 42 °C. *R. toruloides* grown in SCJ had a greater number of down-regulated genes versus up-regulated genes, while growing in 42 °C resulted in the opposite: more genes were up-regulated (**Supplementary Figure S3**). The outermost numbers in the Venn Diagram represent the amount of genes that were exclusively differentially expressed for each condition. The exclusive DEGs can lead to the creation of a new inducible promoter library, given that the promoters will likely be induced by the exact same condition in which their genes were differentially expressed. Although still under development, the current promoter toolbox for *R. toruloides* is limited (Johns, Love and Aves, 2016; Liu *et al*., 2016; Wang *et al*., 2016; Nora, Wehrs, *et al*., 2019). Our dataset can be useful for the development of inducible promoters in the future.

**Figure 2.**
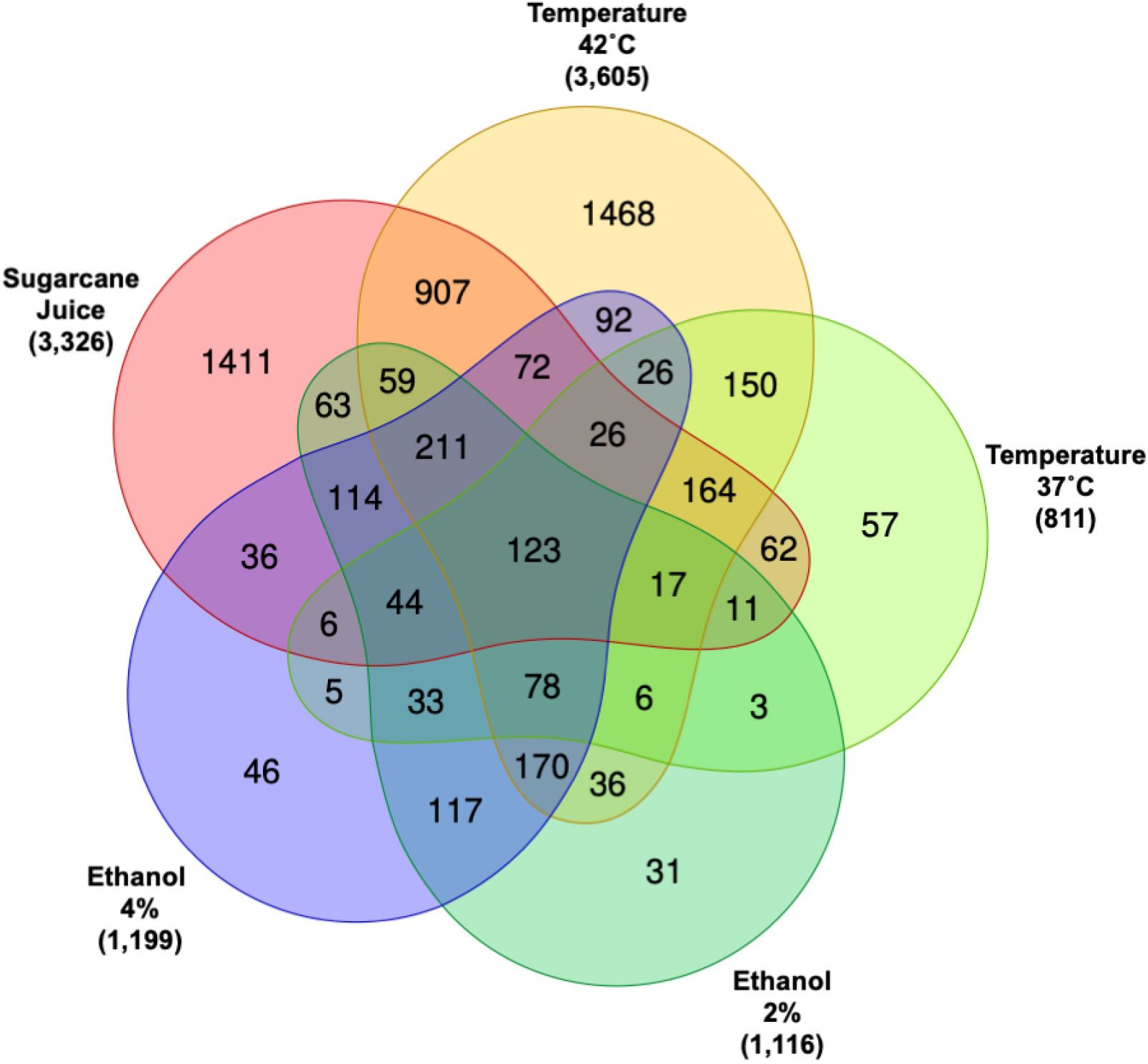
Venn Diagram of DEGs for all conditions. The Venn Diagram shows how many DEGs are shared between all conditions and how many are exclusive for each condition. Numbers outside of the Venn diagram represent the total of DEGs when compared to each respective control.

### 3.3. Functional analyses uncover KEGG pathway enrichment in each condition

To analyze the function of the DEGs, we performed a pathway enrichment analysis. In order to do that, we first had the transcript function annotated by KEGG Orthology. Functional annotation by KEGG was the input required for the R package used for the analysis, called GAGE. Both the annotation and GAGE analysis are described in the methods section. The resulting pathways were filtered to a p-value of less than 0.05. In **Figure 3**, the biochemical pathways of *R. toruloides* that are enriched when using SCJ as a substrate can be seen. The main up-regulated pathways enriched in this condition are those related to cell growth and metabolism, such DNA metabolism and replication, meiosis and amino acid biosynthesis. Pathways related to the biosynthesis of amino acids, ribosomes and cell cycle showed the greatest numbers of up-regulated genes.

**Figure 3.**
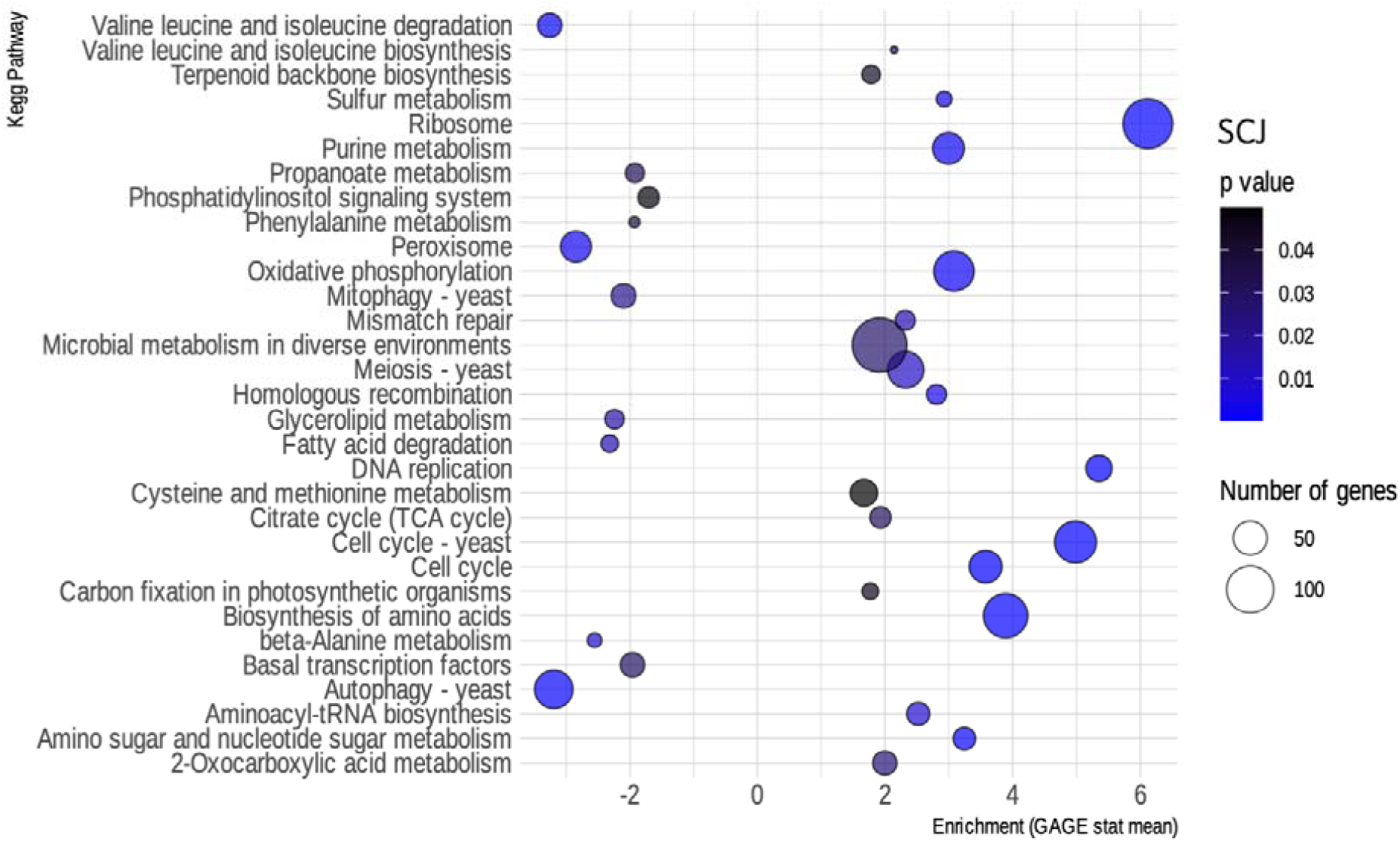
Enriched KEGG pathways for *R. toruloides* grown on SCJ. Bubble map showing the biochemical pathways of *R. toruloides* noted by KEGG enriched in the SCJ condition, as obtained by the GAGE package. Pathways that have an enrichment value greater than 0 are up-regulated, while those that have a value less than 0 are down-regulated. Blue scale inside the bubbles represents the decreasing *p*-values. The different sizes of the bubbles define the approximate number of DEGs in each biochemical pathway.

For the 42 °C condition, although there was a higher number of up-regulated genes (**Supplementary Figure S3**), most of the enriched pathways were found to be down-regulated **(Figure 4)**. Included on that list are several metabolism pathways, showing the exact opposite of what happens in SCJ – metabolism genes are being turned off. In contrast, the condition of 37 °C resulted in a reduced number of differentially expressed transcripts when compared to the control grown at 30 °C (**Supplementary Figure S4**). Yet, in **Supplementary Figure S5** the routes found to be enriched by GAGE by the presence of ethanol in the media can be seen. Most of the pathways are metabolism-related ones that are being down-regulated in both conditions.

**Figure 4.**
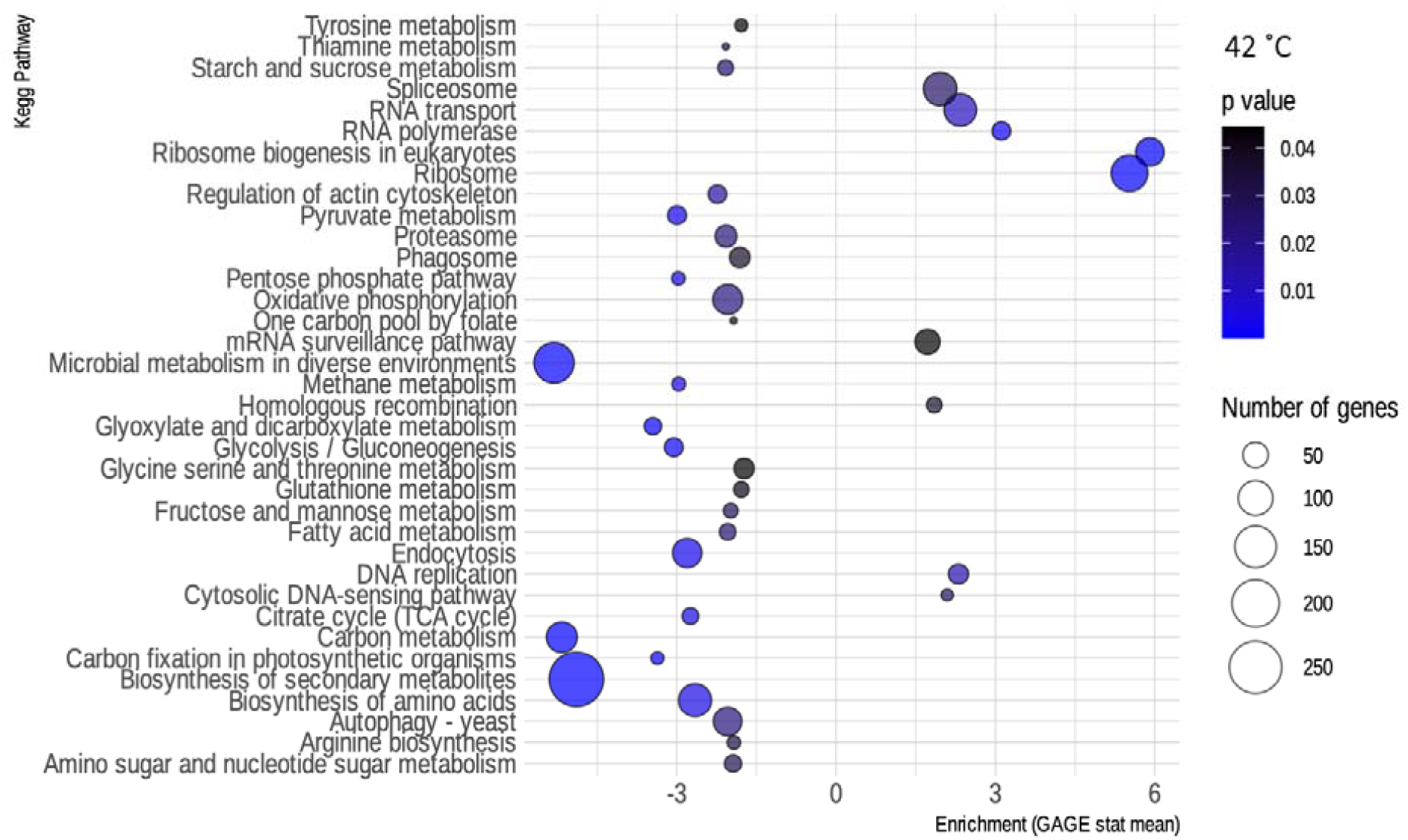
Enriched KEGG pathways for *R. toruloides* grown at 42 °C. Bubble map showing the biochemical pathways of *R. toruloides* noted by KEGG that are enriched in the 42 °C condition, as obtained by the GAGE package. Pathways that have an enrichment value greater than 0 are up-regulated, while those that have a value less than 0 are down-regulated. Blue scale inside the bubbles represents the decreasing p-values. The different sizes of the bubbles define the approximate number of DEGs in each biochemical pathway.

### 3.4. Identification of putative *cis*-regulatory elements and transcription factor candidates

Subsequently, we performed a new functional analysis, this time using the function annotation file of the reference gene created by KOG (provided by the JGI website as described in Methods). The new functional annotation using the KOG database was chosen due to a better correspondence with our transcript IDs resulting in a greater number of categorized genes and less redundancy in gene classes. The most significant classes of genes were selected for further analyses, which can be seen in **Figure 5**. All the remaining classes in which the DEGs of all conditions tested were classified according to KOG annotation are presented in **Supplementary Figure S6**. In both cases, data is shown as the percentage of DEGs in relation to the total number of genes in the *R. toruloides* genome.

**Figure 5.**
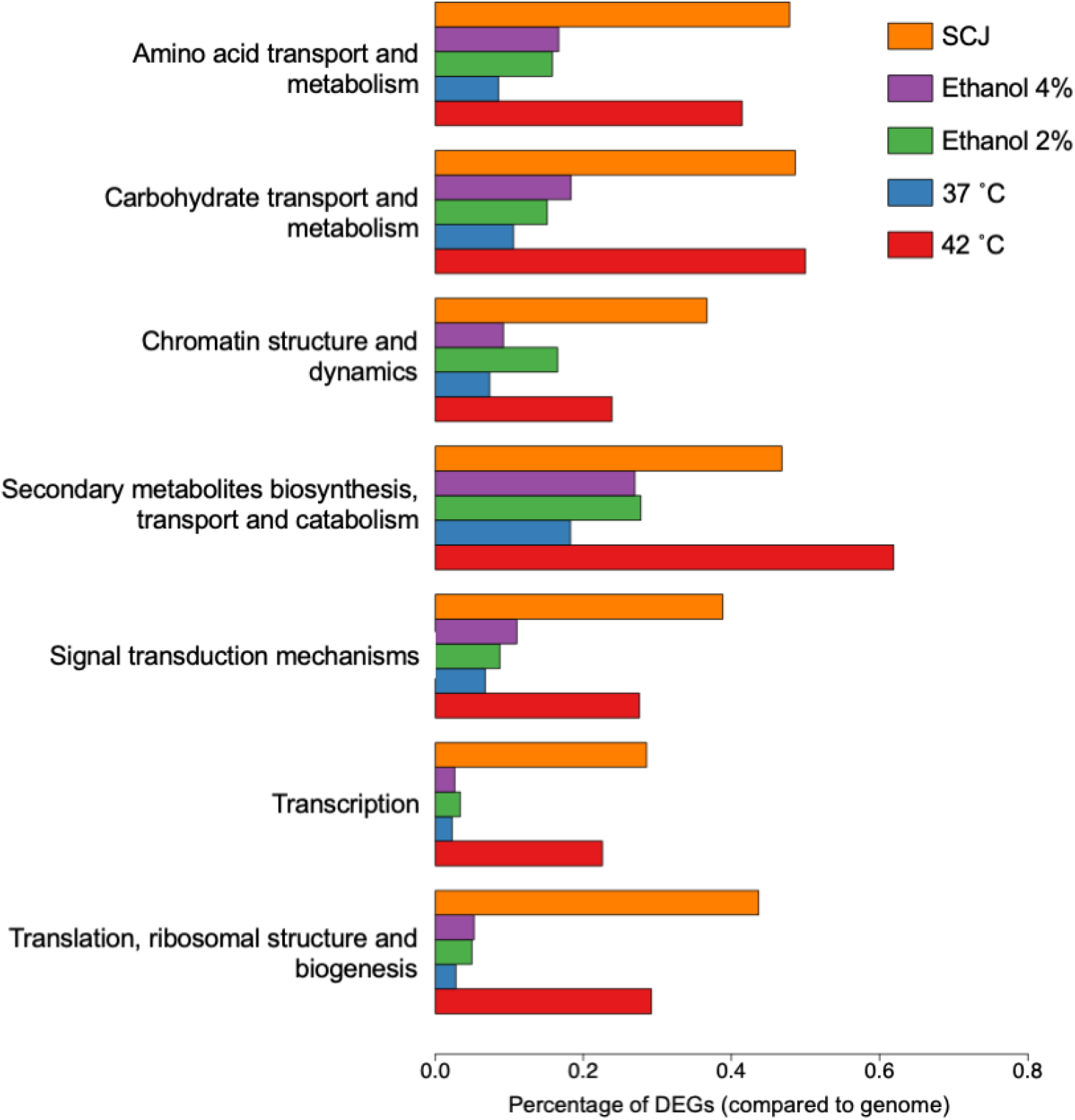
KOG groups selected for TFBS finding. Percentage of DEGs compared to the total number of genes in the *R. toruloides* genome for each condition are shown as annotated using KOG. These are the most relevant conditions with the greatest amount of DEGs and were selected for further analysis. The remaining KOG annotations are shown in **Supplementary Figure 5**.

We used the transcriptomic data generated in this work to search for putative TFs and putative TFBS that might be playing a role in the regulation of both the stress response and the response to complex carbon sources. The pursuit of TF and TFBS was carried out in several stages in order to select suitable candidates. First, we extracted the sequence from the promoters corresponding to the genes of the KOG classes from **Figure 5**. By doing this, we both restricted the quantity of sequences to search motifs, which may reduce background noise, and we attempted to guarantee a biological relationship between these promoter sequences. Then, we used the HOMER software to obtain the putative motifs from those sequences. The motifs obtained were read on the R platform using the universal motif library and a pairwise comparison was performed between all of them, with Pearson’s correlation coefficient (PCC) as a method. The pairwise comparison was done following the rationale that if the motif is found in more than one gene, the chances of being a real motif increase. We also performed pairwise alignment using TOMTOM to complement the analysis. Finally, we filtered the results by a p-value less than 0.05, an e-value less than 1 and a PCC greater than 0.8. The resulting motifs were then compared to the Jaspar fungi 2018 (non-redundant DNA) database using TOMTOM, and some of these alignments can be seen in **Figure 6**. The motif candidates and motifs from JASPAR, aligned by TOMTOM, were plotted in a graph so that we could analyze the similarities between them (**Figure 7)**. The main TFs corresponding to the putative TFBS found using TOMTOM are listed in **Table 2**. The percentage of identity of TF proteins found by this method against proteins from *R. toruloides* genome are also listed in **Table 2**, along with GO functions of the TF proteins.

**Figure 6.**
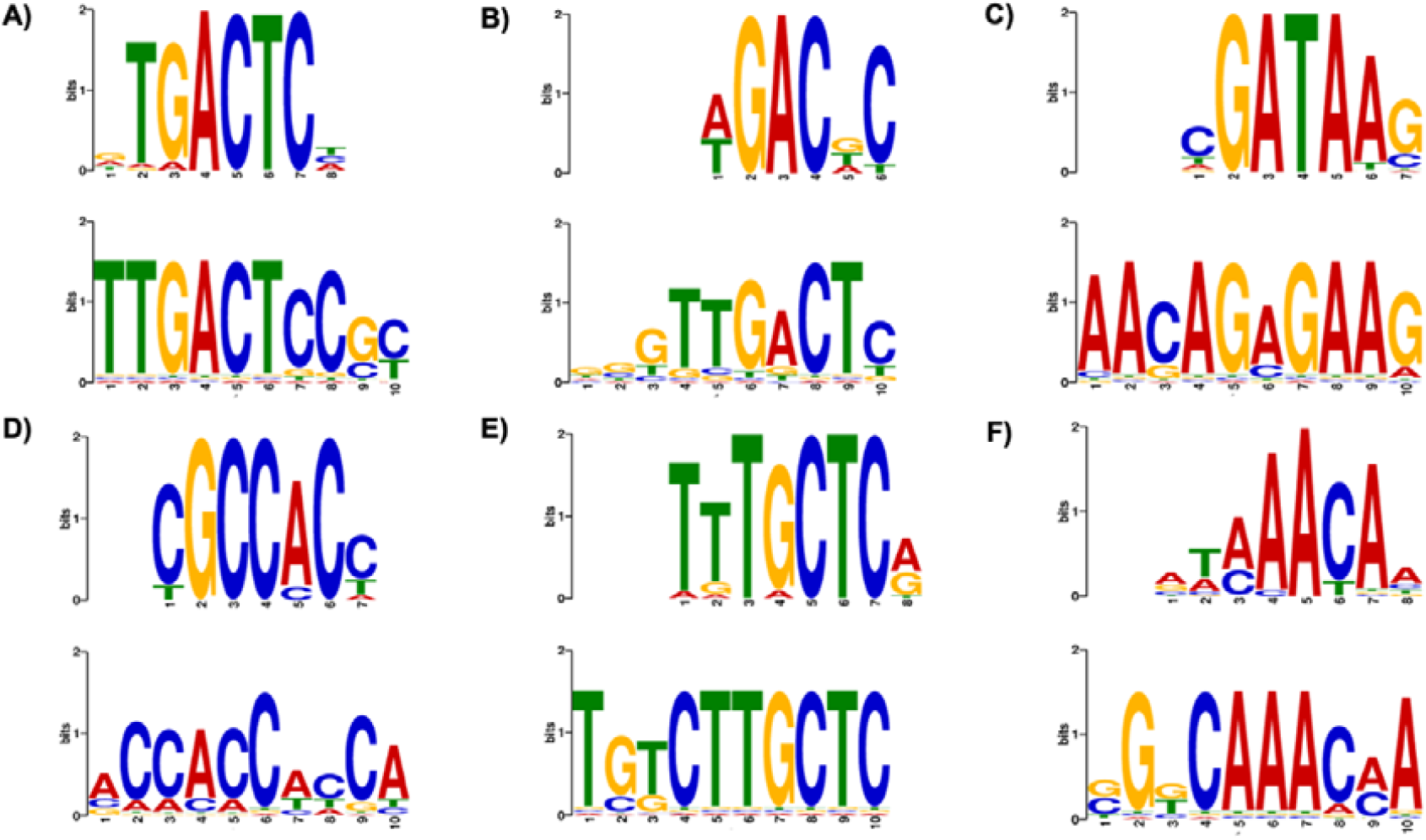
Examples of putative motifs aligned with transcription factors motifs from the JASPAR database using TOMTOM. **(A)** Top motif is the binding motif for ARG81 from *S. cerevisiae*. Bottom motif is from SCJ condition. KOG class: Amino acid transport and metabolism. *p*-value: 1.08e-03. *E*-value: 1.51e+00 **(B)** Top motif for AGR80 from *S. cerevisiae*. Bottom motif is from 42 °C condition. KOG class: Amino acid transport and metabolism. *p*-value: 4.98e-03. *E*-value: 8.76e-01. **(C)** Top motif for DAL80 from *S. cerevisiae*. Bottom motif is from ethanol 2% condition. KOG class: Amino acid transport and metabolism. *p*-value: 3.38e-02. *E*-value: 5.95e+00. **(D)** Top for RPN4 from *S. cerevisiae*. Bottom motif is from 42 °C condition. KOG class: Chromatin structure and dynamics. *p*-value: 4.15e-03. *E*-value: 7.31e-01. **(E)** Top motif for MAC1 from *S. cerevisiae*. Bottom motif is from ethanol 2% condition. KOG class: Secondary metabolites biosynthesis transport and catabolism. *p*-value: 4.15e-03. *E*-value: 7.30e-01. **(F)** Top motif for HCM1 from *S. cerevisiae*. Bottom motif is from SCJ condition. KOG class: Chromatin structure and dynamics. *p*-value: .44e-02. *E*-value: 6.05e+00.

**Figure 7.**
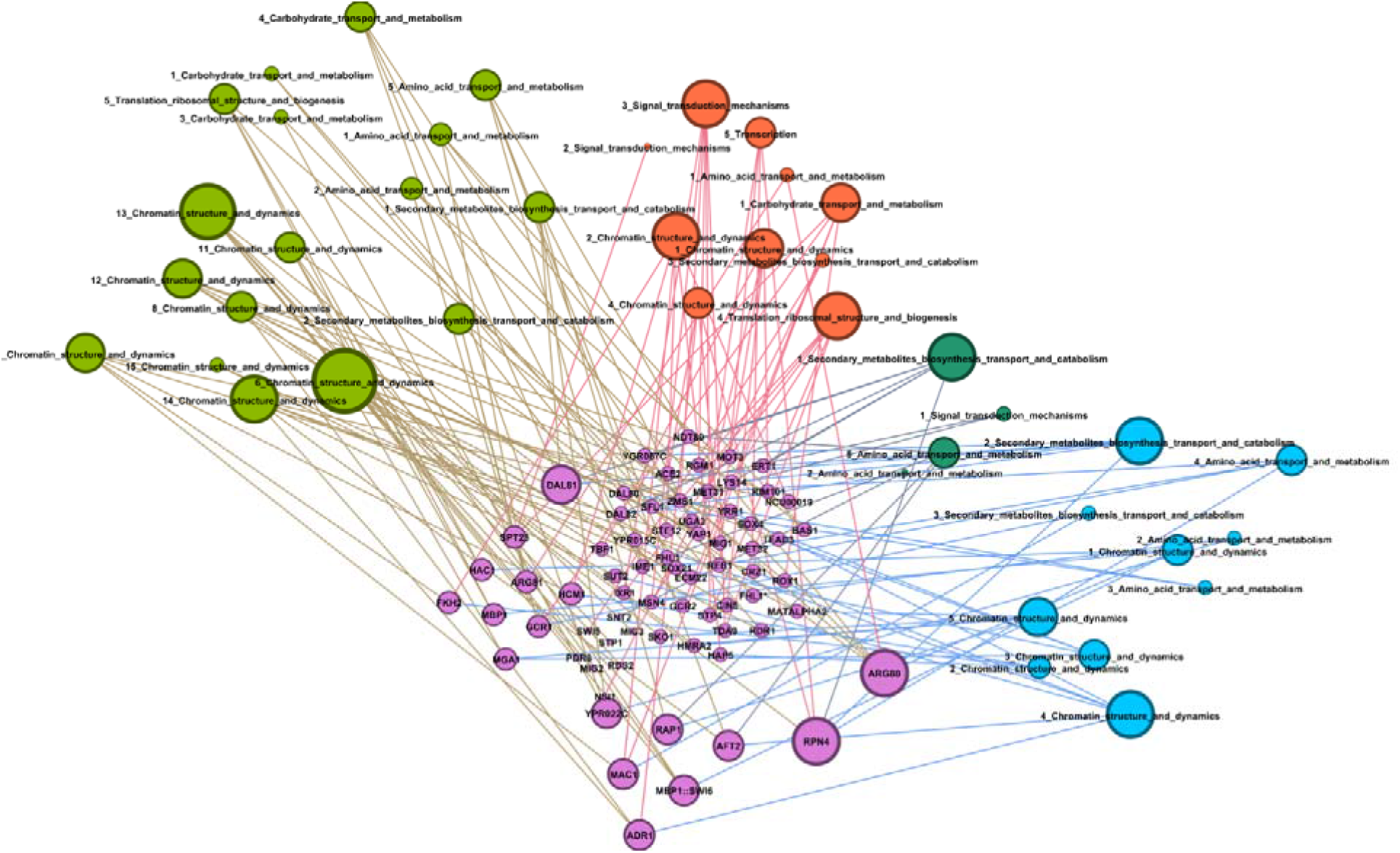
Graph of putative transcription factors and their putative binding sites. Graph showing interaction between putative transcription factors for *R. toruloides* with the TF binding motifs predicted using HOMER. Green nodes represent the sugarcane juice condition, orange nodes represent the 42 °C condition, dark green nodes are the 2% ethanol, and blue nodes are the 4% ethanol condition. Written inside nodes are the number of the TFBS and the name of the gene class annotated by KOG in which that motif was found. Pink nodes are the putative transcription FTs found using TOMTOM.

**Table 2.** Predicted transcription factors and their sequences. BLAST of the predicted transcription factors found by comparison of putative motifs against *R. toruloides* genome showing the percentage of identity for each protein sequence. Protein sequences and Gene Ontology numbers extracted from the Uniprot database. Only some predictive GO functions are shown, according to its correspondence to the study scope.

| TF | GO Number | GO Function | Found in *R. toruloides*? | % of identity |
| --- | --- | --- | --- | --- |
| ADR1 | GO:0061410 | Positive regulation of transcription in response to ethanol | Yes | 54% |
| AFT2 | GO:0034599 | Any process that results in a change in state of a cell as a result of oxidative stress | No | - |
| ARG80 | GO:0006525 | Chemical reactions and pathways involving arginine | Yes | 60% |
| ARG81 | GO:0006525 | Chemical reactions and pathways involving arginine | No | - |
| DAL81 | GO:1901717 | Activates or increases the frequency, rate or extent of gamma-aminobutyric acid catabolic process | No | - |
| FKH2 | GO:0006338 | Dynamic structural changes to eukaryotic chromatin throughout the cell division cycle | Yes | 47% |
| GCR1 | GO:0000433 | Carbon catabolite repression of transcription from RNA polymerase II promoter by glucose | No | - |
| HAC1 | GO:0030968 | Endoplasmic reticulum unfolded protein response | No | - |
| HCM1 | GO:0034599 | Any process that results in a change in state of a cell as a result of oxidative stress | Yes | 49% |
| MAC1 | GO:0045732 | Positive regulation of protein catabolic process | Yes | 50% |
| MBP1 | GO:0071931 | Positive regulation of transcription involved in G1/S transition of mitotic cell cycle | Yes | 52% |
| MGA1 | GO:0007124 | Pseudohyphal growth in response to an environmental stimulus | Yes | 51% |
| RAP1 | GO:0010833 | Telomere maintenance via telomere lengthening | No | - |
| RPN4 | GO:0097201 | Negative regulation of transcription from RNA polymerase II promoter in response to stress | No | - |
| SPT23 | GO:0070417 | Cellular response to cold | Yes | 45% |
| SWI6 | GO:0061408 | Positive regulation of transcription from RNA polymerase II promoter in response to heat stress | Yes | 39% |
| YPR022C | GO:0000981 | Modulates the transcription of specific gene sets transcribed by RNA polymerase II | Yes | 69% |

## 4. Discussion

In this work, we developed a reproducible bioinformatics workflow that can be used for the discovery of regulatory elements using transcriptomic data as input, and applied to the emergent microbial cell factory *R. toruloides*. First, by analyzing the transcriptomic data, we showed that *R. toruloides* presented different transcriptional responses to all the conditions tested. When using media containing SCJ and urea, functional analysis showed a predominance of metabolism pathways being up-regulated, while degradation pathways such as autophagy and peroxisome are down-regulated. This pattern is reasonable since the yeast adapted its metabolism from the inoculum containing no sugar to the rich and complex environment provided by SCJ. Interestingly enough, one of the pathways that were found to be up-regulated by GAGE was the terpenoid backbone biosynthesis. This is a good indicator that we can use the abundant and inexpensive media composed by sugarcane and urea as a start point substrate to produce terpenes in *R. toruloides*, like previously done using corn stover hydrolysates (Kirby *et al*., 2021). Moreover, when growing the yeast at 42 °C, cytosolic DNA sensing pathways and mRNA surveillance pathways were found to be up-regulated and are representative of cell stress responses (Jamar, Kritsiligkou and Grant, 2017; Meng *et al*., 2021). Similarly, in this condition, the homologous recombination pathway is being up-regulated – possibly for dealing with cell damaging (Litwin *et al*., 2013). We also hypothesized that the transcriptional responses to the two high temperature conditions can be classified into two distinct types of stress responses. The 42 °C condition caused the heat-shock response, when cells repress protein biosynthesis – as we can see by the decrease of the biosynthesis of amino acids - and protein misfolding and oxidative stress occur (Verghese *et al*., 2012). In contrast, the condition of 37 °C might have reached the other type of response: thermotolerance – given that most transcripts were not differentially expressed compared to the control grown at 30 °C. The very few pathways found to be enriched in this condition, like cell cycle and oxidative phosphorylation, might suggest that *R. toruloides* is enduring the increase in temperature (Shui *et al*., 2015; Huang *et al*., 2018). Nevertheless, another study showed that cultivating *R. toruloides* at 37 °C greatly impaired its growth (Wu *et al*., 2020). Yet, the ethanol response greatly corresponds with what is described in literature for ethanol stress and tolerance in *S. cerevisiae* (Stanley *et al*., 2010). It is interesting to note that cell cycle and DNA replication pathways are both being down-regulated, showing inhibition of cell growth, as it was expected for ethanol stress (Stanley *et al*., 2010). Inhibition of endocytosis was also reported as an ethanol stress response (Lucero *et al*., 2000) and we can see that both concentrations of ethanol resulted in a decrease of expression of *R. toruloides* endocytosis genes. Additionally, the proteasome genes are up-regulated, demonstrating that cells are dealing with misfolded proteins, which is described in literature as a sign of ethanol stress (Bubis *et al*., 2020). Hence, we hypothesized that the response to the two different concentrations of ethanol is still a stress response and *R. toruloides* has not yet been able to achieve tolerance to the compound (Lucero *et al*., 2000; Stanley *et al*., 2010; Bubis *et al*., 2020).

Furthermore, we explored the cis-regulatory elements related to the DEGs found in the transcriptomic analysis. The TFBS predicted for *R. toruloides* presented similarity mainly with binding sites for the following TFs: ARG80, RPN4, ADR1 and DAL81, and those TFs can be seen in the graph **(Figure 7)** as having the greater number of similar putative motifs. As demonstrated in **Table 2**, HAC1 protein is involved in ER-unfolded protein response and RPN4 protein is involved in response to stress, which could explain the enrichment of both their binding motifs in our industrial stress conditions. In fact, studies showed that mutations in RPN4 protein conferred ethanol resistance to *S. cerevisiae* (Bubis *et al*., 2020). However, there were no homologs of these proteins found in our carotenogenic yeast genome. Some TFs were found to be responsive to oxidative stress, such as AFT2 and HCM1, whereas only HCM1 was found in the *R. toruloides* genome. Interestingly, DAL81 protein was found to be mutated in response to high temperature stress in a study by Huang *et al*. (2018). This could explain the enrichment of its binding motifs in our heat shock transcripts, although a homolog of this protein was not found. SWI6 binding motifs may be an interesting source for future studies, since this protein was found in *R. toruloides* genome with 39% identity and it is a protein that regulates heat stress response. Interestingly, binding motifs for SPT23, a protein that regulates cold response, were also found. This protein was found in the genome of *R. toruloides* with 45% identity. Nevertheless, the binding of the TF proteins to their respective motifs discovered by our pipeline needs to be further tested to confirm their activity. Alternatively, in the case of the ones whose corresponding binding protein homologs were not found, recognition of the identified motifs could be performed by yet unknown TFs. As shown by Antonieto *et al*., (2019), a transcriptional regulator with low homology to AZF1 recognizes a well-conserved motif regulating expression of cellulases genes.

The motifs found can be used as starting points for new transcriptional modulation studies. The ultimate goal would be to define a core promoter sequence for *R. toruloides*, and then change the sequences of TFBS according to the desired behavior. For instance, one can study ethanol tolerance in this organism by adding the RPN4 consensus sequence to their respective constructs. Additionally, our pipeline can be applied to other organisms that already have well-defined core promoter sequences, where scientists can “play around” with both motif sequences and locations in order to unlock new transcriptional behaviors (Monteiro *et al*., 2020; Tominaga, Kondo and Ishii, 2022). Also, as mentioned in the Results section, our transcriptomic data can be used to create new promoter libraries for our yeast. Ultimately, anyone can resort to our tools to generate better transcriptional understanding of their organism of interest.

## 5. Conclusion

Here, we characterized the transcriptional responses of *R. toruloides* when growing in industry-like conditions, and discovered new regulatory elements that were enriched in each context. We showed how differential gene expression, followed by a custom, reproducible bioinformatics motif discovery workflow, was able to predict putative motifs for binding of transcription factors in our strain, most of them involved in stress-related responses. Using our approach, DNA motifs similar to the binding sites for ADR1, ARG80, DAL81 and RPN4 transcription factors were found to be the most abundantly enriched in our dataset. While ADR1 and ARG80 proteins were found in *R. toruloides* genome with 54% and 60% of identity, respectively, DAL81 and RPN4 were not found. The novel putative *cis*-regulatory elements described here offer a great initial point to optimize gene regulation in industrial conditions expanding the current knowledge on regulatory networks for this yeast. Furthermore, our pipeline for motif discovery can be easily applied for other unexplored hosts.

## Data availability statement

Raw data is available as Sequencing Read Archives (SRA) on the NCBI website under accession number PRJNA883675 (https://www.ncbi.nlm.nih.gov/sra/PRJNA883675).

## Author contributions

RSR, MEG, LCN, and MHAC designed the project. LCN and IPS performed RNA experiments. MHAC, LCN and IPS performed the differential expression analysis. MHAC developed the motif discovery pipeline. RRS contributed substantially to the analysis, interpretation of data, and availability of the developed workflow. LCN wrote the manuscript. All authors critically revised the manuscript and approved the submitted version.

## Funding

This work was supported by the São Paulo Research Foundation (FAPESP), the National Council for Scientific and Technological Development (CNPq), and CAPES, all from Brazil. LCN was funded by FAPESP grant number 2019/04942-7 and CNPq grant number 140212/2019-1. M-EG was funded by FAPESP grant number 2021/01748-5. RSR was funded by FAPESP grant numbers 2019/15675-0. MHAC and IPS were funded by CAPES. RRS was funded by FAPESP grant numbers 2017/18922-2, 2019/05026-4 and 2020/02207-5.

## Conflict of interest

The authors declare that the research was conducted in the absence of any commercial or financial relationships that could be construed as a potential conflict of interest.

## Abbreviations

DEG: differentially expressed genes
GO: Gene Ontology
KEGG: Kyoto Encyclopedia of Genes and Genomes
KOG: EuKaryotic Orthologous Groups
PCC: Pearson’s correlation coefficient
SCJ: sugarcane juice
TF: transcription factor
TFBS: transcription factor binding site

## Notes

https://www.ncbi.nlm.nih.gov/sra/PRJNA883675

https://mycocosm.jgi.doe.gov/Rhoto_IFO0880_4/Rhoto_IFO0880_4.home.html

https://www.genome.jp/tools/kaas/

http://uniprot.org

https://github.com/computational-chemical-biology/cis_reg

### Competing Interest Statement

The authors have declared no competing interest.

