## Supplementary Material for "Mining novel cis-regulatory elements from the emergent host *Rhodosporidium toruloides* using transcriptomic data"

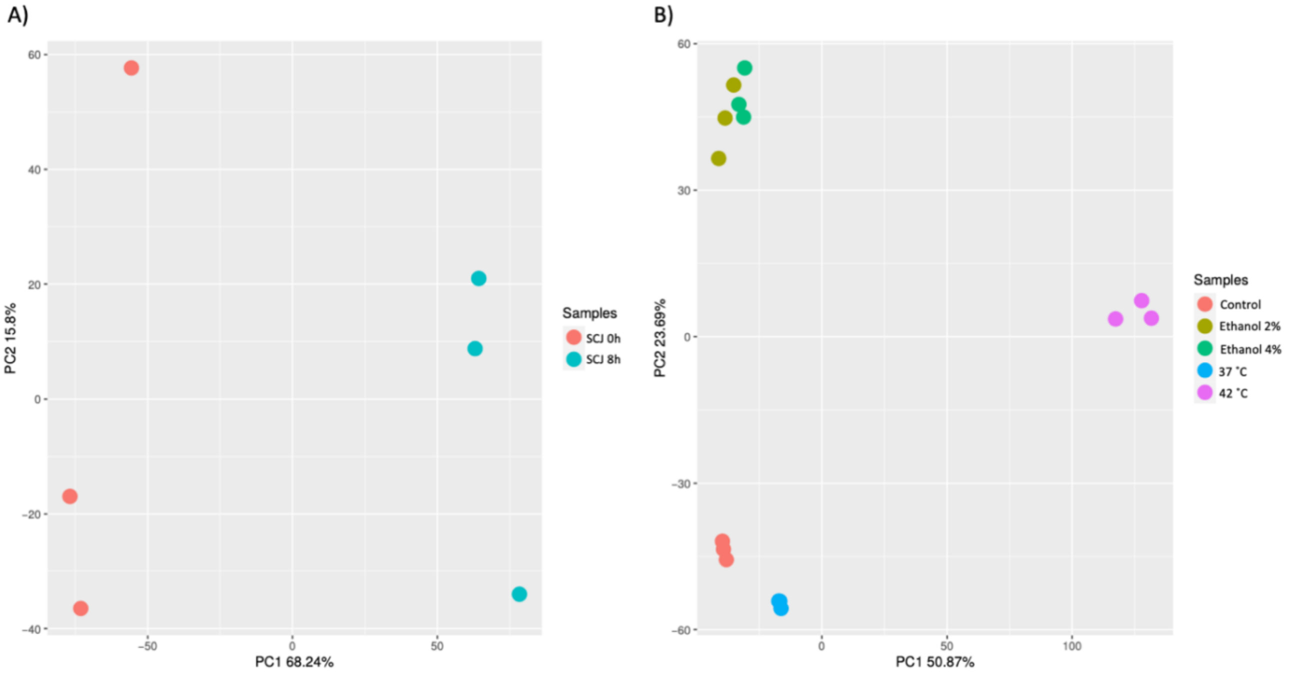


**Supplementary Figure S1. PCA of experimental replicates.** Principal Component Analysis (PCA) applied to *R. toruloides* transcripts, grouping the conditions based on the similarity of the projection. **(A)** Experiment using SCJ as a substrate. The conditions of this experiment are time 0 h (control cultures grown in LB for 24 hours) and 8 h (cultures transferred from the pre-inoculum to a medium containing sugarcane juice and urea). **(B)** Experiment using industrial stress conditions. The conditions for this experiment are control time (grown in YPD for 24 hours) and the respective stress conditions: 2% ethanol, 4% ethanol, 42 ˚C and 37 ˚C.

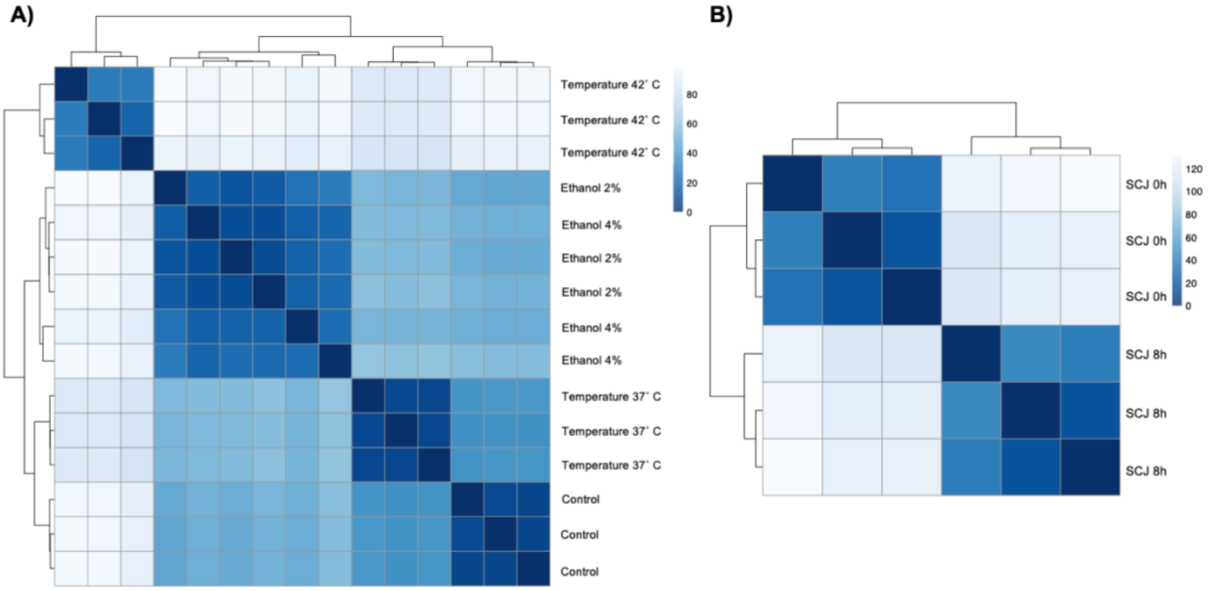


**Supplementary Figure S2. Heatmap for experimental replicates measured by Euclidean Distance. (A)** Heatmap showing the correlation of gene expression between transcripts of *R. toruloides* grown under conditions of industrial stress. The conditions for this experiment are control (grown in YPD for 24 hours) and the respective stress conditions: ethanol 2%, ethanol 4%, 42 ˚C and 37 ˚C. **(B)** Heatmap showing the correlation of gene expression between transcripts of *R. toruloides* grown in SCJ. The scale represents Euclidean distance, where the lighter the blue, the greater the distance between the samples.

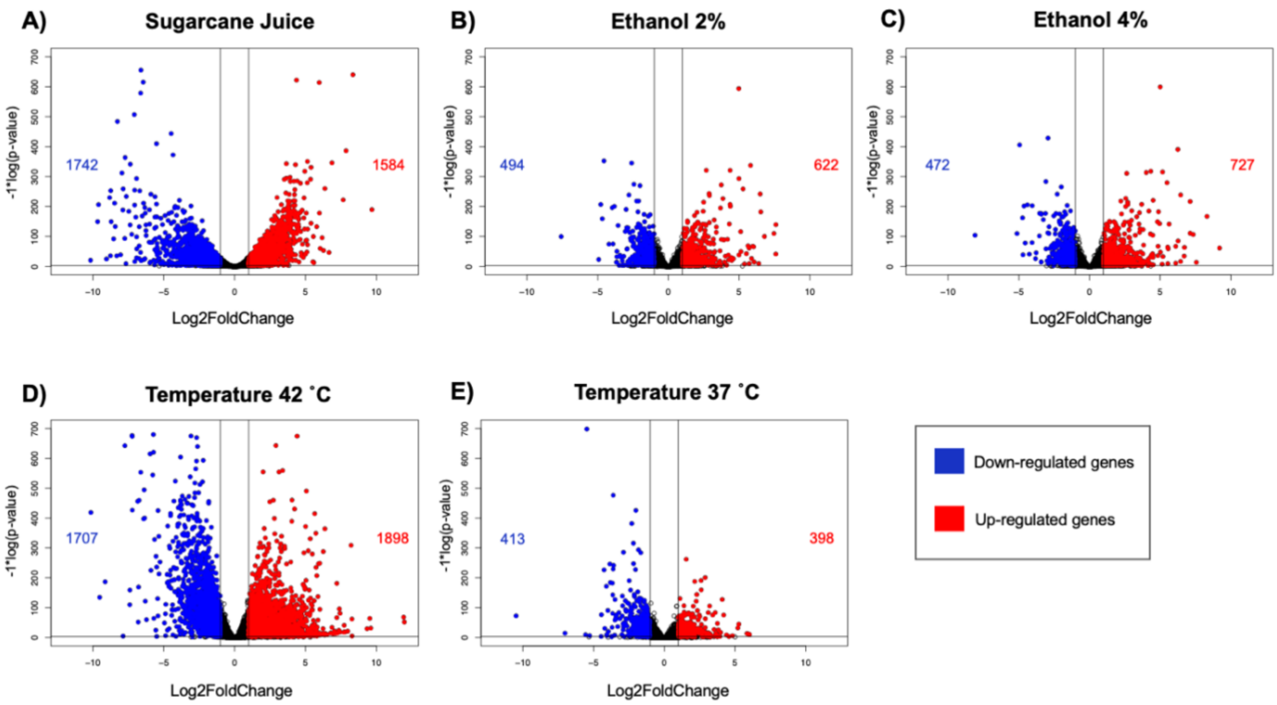


**Supplementary Figure S3. Volcano plots for *R. toruloides* DEGs in all conditions.** Volcano plot showing the DEGs for each experimental setting when compared to its respective control. Genes that are down-regulated are represented in blue and genes that are up-regulated are represented in red. The vertical cut lines on the graph divide the log2FoldChange values less than -1 (left) and greater than 1 (right). The horizontal cut line in the graph divides the valid p-value values less than 0.05 in -log (*p*-value).

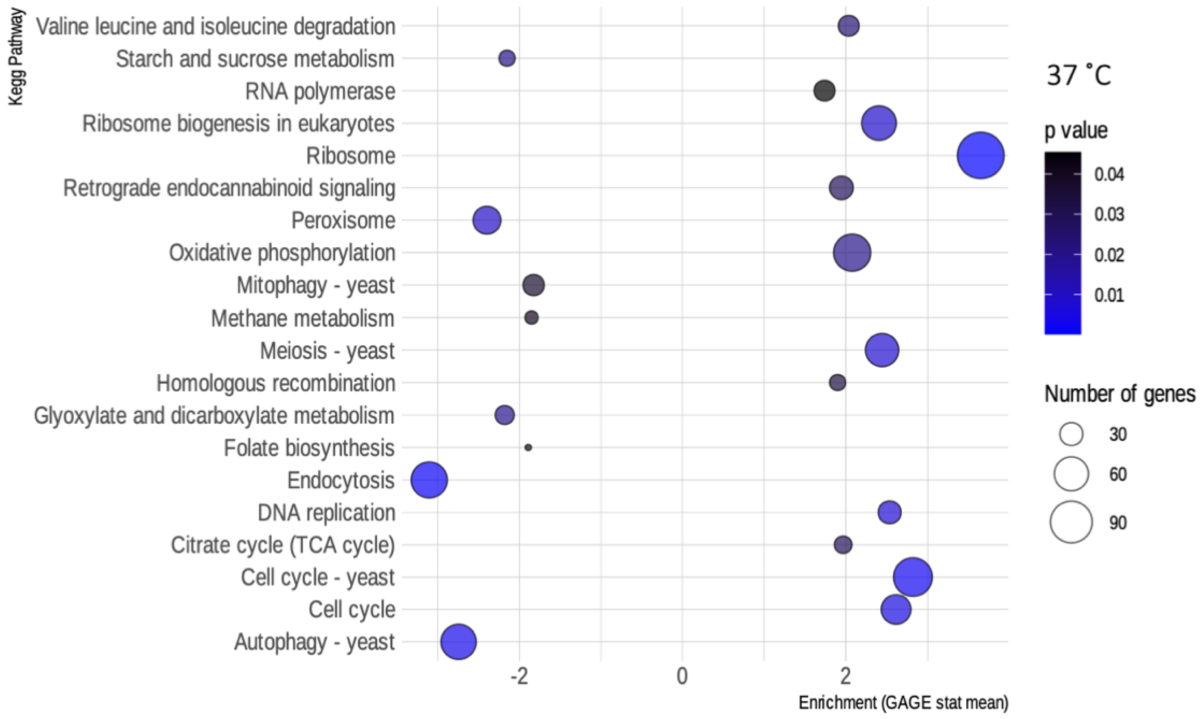


**Supplementary Figure S4. Enriched KEGG pathways for *R. toruloides* grown at 37 ˚C.** Bubble map showing the biochemical pathways of *R. toruloides* noted by KEGG that are enriched in the 37 ˚C condition, as obtained by the GAGE package. Pathways that have an enrichment value greater than 0 are up-regulated while those that have a value less than 0 are down-regulated. Blue scale inside the bubbles represents the decreasing *p*-values. The different sizes of the bubbles define the approximate number of DEGs in each biochemical pathway.

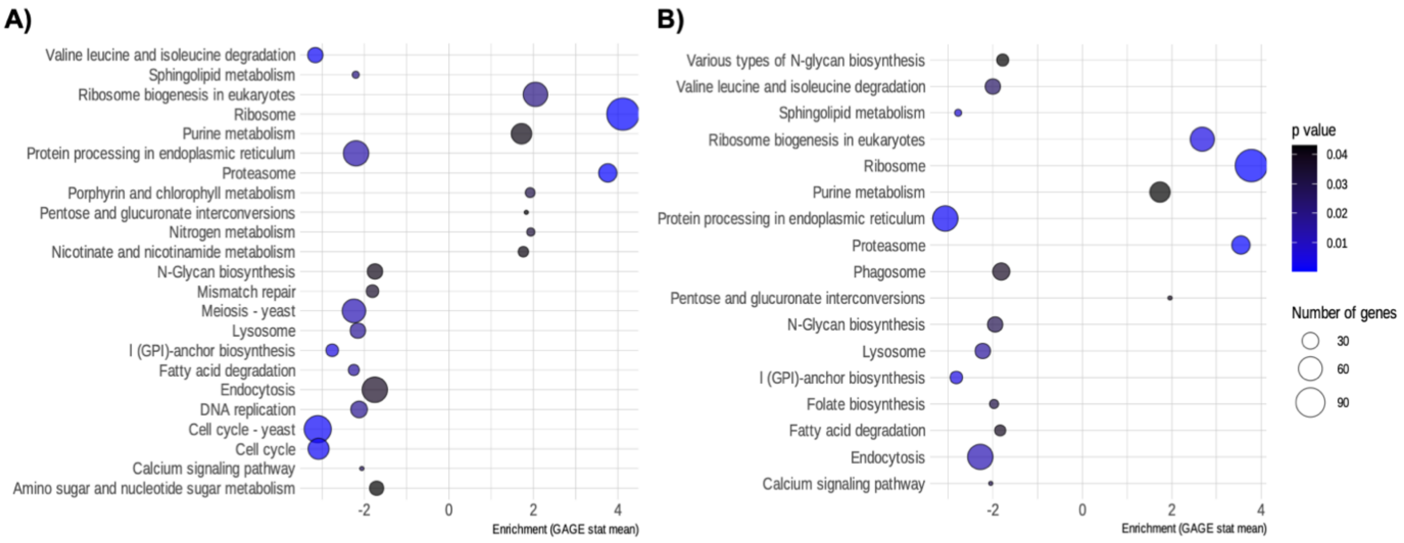


**Supplementary Figure S5. Enriched KEGG pathways for *R. toruloides* grown in ethanol conditions.** Bubble map showing the biochemical pathways of *R. toruloides* noted by KEGG that are enriched in the ethanol conditions, as obtained by the GAGE package. **(A)** Ethanol 2%. **(B)** Ethanol 4%. Pathways that have an enrichment value greater than 0 are up-regulated while those that have a value less than 0 are down-regulated. Blue scale inside the bubbles represents the decreasing *p*-values. The different sizes of the bubbles define the approximate number of DEGs in each biochemical pathway.

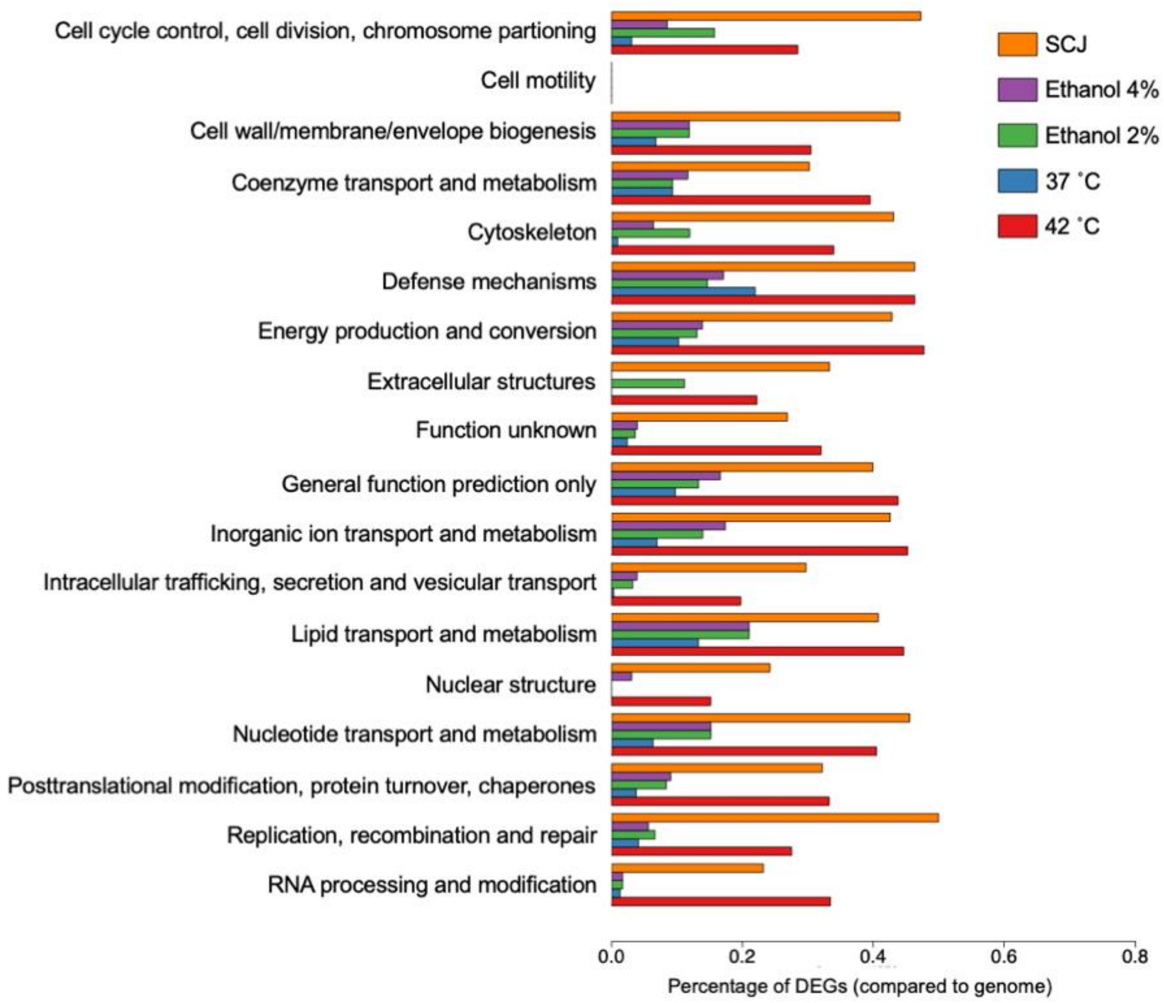


**Supplementary Figure S6. Percentage of DEGs annotated using KOG compared to the total number of genes in the *R. toruloides* genome.** Percentage of DEGs compared to the total number of genes in the *R. toruloides* genome for each condition are shown as annotated using KOG.
